# *Mcmdc2*, a minichromosome maintenance (MCM) paralog, is required for repair of meiotic DNA breaks and homolog pairing in mouse meiosis

**DOI:** 10.1101/080887

**Authors:** Adrian J. McNairn, Vera D. Rinaldi, John C. Schimenti

**Affiliations:** Cornell University, College of Veterinary Medicine, Dept. of Biomedical Sciences, Ithaca NY 14853 USA

**Author notes:** John Schimenti, PhD, Cornell University, College of Veterinary Medicine, Ithaca, NY 14853.

**Keywords:** meiosis, recombination, mouse, double strand break repair, synapsis

## Abstract

The mammalian *Mcm-domain containing 2* (*Mcmdc2*) gene encodes a protein of unknown function that is homologous to the mini-chromosome maintenance family of DNA replication licensing and helicase factors. *Drosophila melanogaster* contains two separate genes, the “Mei-MCMs,” that appear to have arisen from a single ancestral *Mcmdc2* gene. The Mei-MCMs are involved in promoting meiotic crossovers by blocking the anti-crossover activity of BLM helicase, a function performed by MSH4 and MSH5 in metazoans. Here, we report that MCMDC2-deficient mice of both sexes are viable but sterile. Males fail to produce spermatozoa, and formation of primordial follicles is disrupted in females. Histology and immunocytological analyses of mutant testes revealed that meiosis is arrested in Prophase I, and is characterized by persistent meiotic double-stranded DNA breaks (DSBs), failure of homologous chromosome synapsis and XY body formation, and an absence of crossing over. These phenotypes essentially phenocopy those of MSH4/5 deficient meiocytes. The data indicate that MCMDC2 is essential for invasion of homologous sequences by RAD51- and DMC1-coated ssDNA filaments, or stabilization of recombination intermediates following strand invasion, both of which are needed to drive stable homolog pairing and DSB repair via recombination in mice.

## Introduction

The MCM family of proteins were discovered based on their crucial functions in DNA replication, and contain conserved “MCM” and ATPase domains (Tye 1999). However, there are additional MCM family members that function outside of the core MCM2-7 replicative helicase complex. MCM8 and MCM9 function in DNA repair and homologous recombination (Park *et al.* 2013; Traver *et al.* 2015). Both are dispensable for DNA replication in mice, but MCM8 and MCM9-deficient cells exhibit defects in homologous recombination repair in response to DNA damage (Hartford *et al.* 2011; Lutzmann *et al.* 2012; Nishimura *et al.* 2012; Park *et al.* 2013; Lee *et al.* 2015; Luo and Schimenti 2015). Surprisingly *Mcm9*, which is absent from *Drosophila*, is also required for DNA mismatch repair, and this may actually be its primary function (Traver *et al.* 2015). *Mcm8* null mice of both sexes are sterile due to defects in homologous recombination repair during meiotic prophase I (Lutzmann *et al.* 2012), whereas *Mcm9* mutant mice are defective in primordial germ cell proliferation that leads to reduced (males) or absent (females) germ cells (Lutzmann *et al.* 2012) (Hartford *et al.* 2011).

During meiosis in many organisms including mice, homologous chromosome pairing and synapsis is driven by recombination, and proper segregation of homologs during the first meiotic division depends upon each chromosome pair undergoing at least 1 crossover (CO) event. To produce COs, there is an orchestrated process in which abundant programmed DNA double stranded breaks (DSBs; ~200/cell in mice) are produced, followed by repair of the majority (~90%) of these breaks by non-crossover (NCO) repair, and the remainder via CO recombination (Handel and Schimenti 2010). The mechanisms regulating CO vs NCO recombination is an area of intense study. Most of the CO events in both yeast and mice require proteins of the ZMM complex, including MSH4 and MSH5, which stabilize recombination intermediates and facilitate double Holliday junction formation after initial D-loops are formed by invasion of a single stranded end of a resected DSB (Manhart and Alani 2016). In part, this is done by inhibiting the anti-CO activity of BLM helicase, which otherwise promotes synthesis-dependent strand annealing (SDSA) and NCO repair (Holloway *et al.* 2010; De Muyt *et al.* 2012). In *Drosophila*, which lacks MSH4 and MSH5, a complex consisting of MEI-217, MEI-218, and REC (the latter being an MCM8 ortholog), termed Mei-MCM (meiotic minichromosome maintenance), assumes the role of promoting COs by inhibiting BLM helicase (Kohl *et al.* 2012).

BLM is a conserved member of RecQ family of DNA helicase that was identified as a rare recessive genetic disorder in humans (Bloom’s Syndrome), characterized by dwarfism, increased cancer susceptibility, and immune deficiency. Mutations in the *Drosophila Blm* gene result in female sterility, defects in DNA repair, and increased sensitivity to ionizing radiation (Adams *et al.* 2003; Mcvey *et al.* 2007). In mice, *Blm* deficiency causes embryonic lethality, though heterozygosity or conditional somatic deletion causes increased genomic instability and tumor susceptibility (Chester *et al.* 1998; Luo *et al.* 2000; Goss *et al.* 2002; Chester *et al.* 2006). Conditional germline deletion of *Blm* in mice disrupts meiotic prophase I progression; mutant spermatocytes exhibited aberrations in chromosome synapsis and elevated numbers of COs (Holloway *et al.* 2010).

The Mei-MCM complex in *Drosophila* was suggested to have evolved to assume the anti-CO activities of MSH4/MSH5. Interestingly, MEI-217 and MEI-218 seem to have arisen from a single ancestral MCM (minichromosome maintenance) gene, in that the former contains an MCM N-terminal domain and the latter a C-terminal AAA ATPase domain, and these two genes are adjacent in the genome and are expressed as a bicistronic transcript (Kohl *et al.* 2012) (Liu *et al.* 2000). The mammalian ortholog is a single gene called *Mcmdc2* (Mcm domain containing protein 2) that resembles a typical MCM protein with the aforementioned domain structure. However, unlike other members of the MCM family, *Mcmdc2* encodes a much smaller MCM-domain than that which is present in other MCM proteins, and the ATPase domains of both MCMDC2 and Mei-218 contain amino acid changes predicted to disrupt actual ATPase activity (Kohl *et al.* 2012). Mutation of any of the Mei-MCMs (*rec*, *mei-217* or *mei-218*) in *Drosophila* causes a severe reduction of COs but not NCO recombination (gene conversion), leading to increased non-disjunction and reduced fertility (Blanton *et al.* 2005) (Grell 1984) (Kohl *et al.* 2012) (Liu *et al.* 2000).

*Mcmdc2* mRNA is present most abundantly in mouse testis (e.g. EMBL-EBI Expression atlas; GEO datasets GSE43717, GSE39970, GSE44346). Though frequently amplified in various types of cancer (cBioPortal), this gene is essentially unstudied in mammals. To explore the potential role of mammalian *Mcmdc2*, and explore whether it performs a role similar to the *Drosophila mei-217/218* and/or mammalian *Msh4/Msh5*, we generated mice bearing a mutant allele. Phenotypic and immunocytological analyses revealed that it is crucial for repair of meiotic DSBs, chromosome synapsis and fertility, but not other discernable somatic processes. The phenotype is similar to that of MSH4/5, suggesting that it may be involved in aiding their roles in stabilizing recombination intermediates and promoting interhomolog recombination and crossovers.

## Results and Discussion

To determine the function of mammalian *Mcmdc2*, we generated knockout mice from ES cells. The targeted allele was of a gene trap design, whereby the normal transcript is truncated by virtue of an introduced splice acceptor that created a lacZ fusion. Additionally, a neomycin selectable marker flanked by loxP sites was integrated into an intron (Fig. S1A). This allele is referred to as *Mcmdc2^Gt^*. Founder animals were genotyped by PCR and the expression pattern of *Mcmdc2* was analyzed by RT-qPCR and lacZ activity staining (Figs. S1B; S2). These analyses showed that *Mcmdc2* is expressed primarily in gonads and brain, but not in heart, lung, or in MEFs (mouse embryonic fibroblasts). These results are consistent with data from transcriptome datasets as mentioned in the Introduction.

While all 6 genes encoding subunits of the MCM replicative helicase (*Mcm2-7*) are essential for viability, *Mcm8* and *Mcm9* are not (Hartford *et al.* 2011; Lutzmann *et al.* 2012; Luo and Schimenti 2015). To determine if *Mcmdc2* is also essential, *Mcmdc2^Gt/+^* heterozygotes were intercrossed and the offspring genotyped. Consistent with the limited range of expression, all genotypes were obtained in Mendelian ratios, indicating the gene is not required for viability (χ^2^ = 0.47; Table 1). Homozygotes appeared phenotypically normal, but both male and female animals were found to be sterile when bred to WT animals.

**Table 1.** *Mcmdc2^Gt/Gt^* mice are viable. Numbers were produced from an intercross of heterozygotes.

|  | **+/+** | ***Mcmdc2*<sup>Gt/+</sup>** | ***Mcmdc2*<sup>Gt/Gt</sup>** |
| --- | --- | --- | --- |
| **Male** | 17 | 36 | 21 |
| **Female** | 14 | 41 | 20 |
| **Total** | 31 | 77 | 41 |

To identify the cause of sterility, we examined gonads of mutant mice. Individual testes isolated from 12-week old males were much smaller than WT or heterozygous littermates of the same age (Fig. 1A, B), weighing ~4 times less than those of littermates (0.031g ± 0.0013 versus 0.13g ± 0.005) (Fig 1B). Histological analyses of mutant testes showed a lack of postmeiotic spermatids, resulting in seminiferous tubules having luminal areas devoid of normal germ cells compared to heterozygous or WT littermates (Fig. 1C). Similar to most meiotic mutant mice defective in recombination or chromosome synapsis, spermatocytes were arrested in a developmental stage resembling pachynema (Handel and Schimenti 2010). Histology of mutant ovaries revealed an absence of primordial or more advanced follicles in both adult and wean age (24 dpp) females (Fig. 2A,B). Therefore, disruption of *Mcmdc2* causes catastrophic failure of gametogenesis in both sexes.

**Figure 1.**
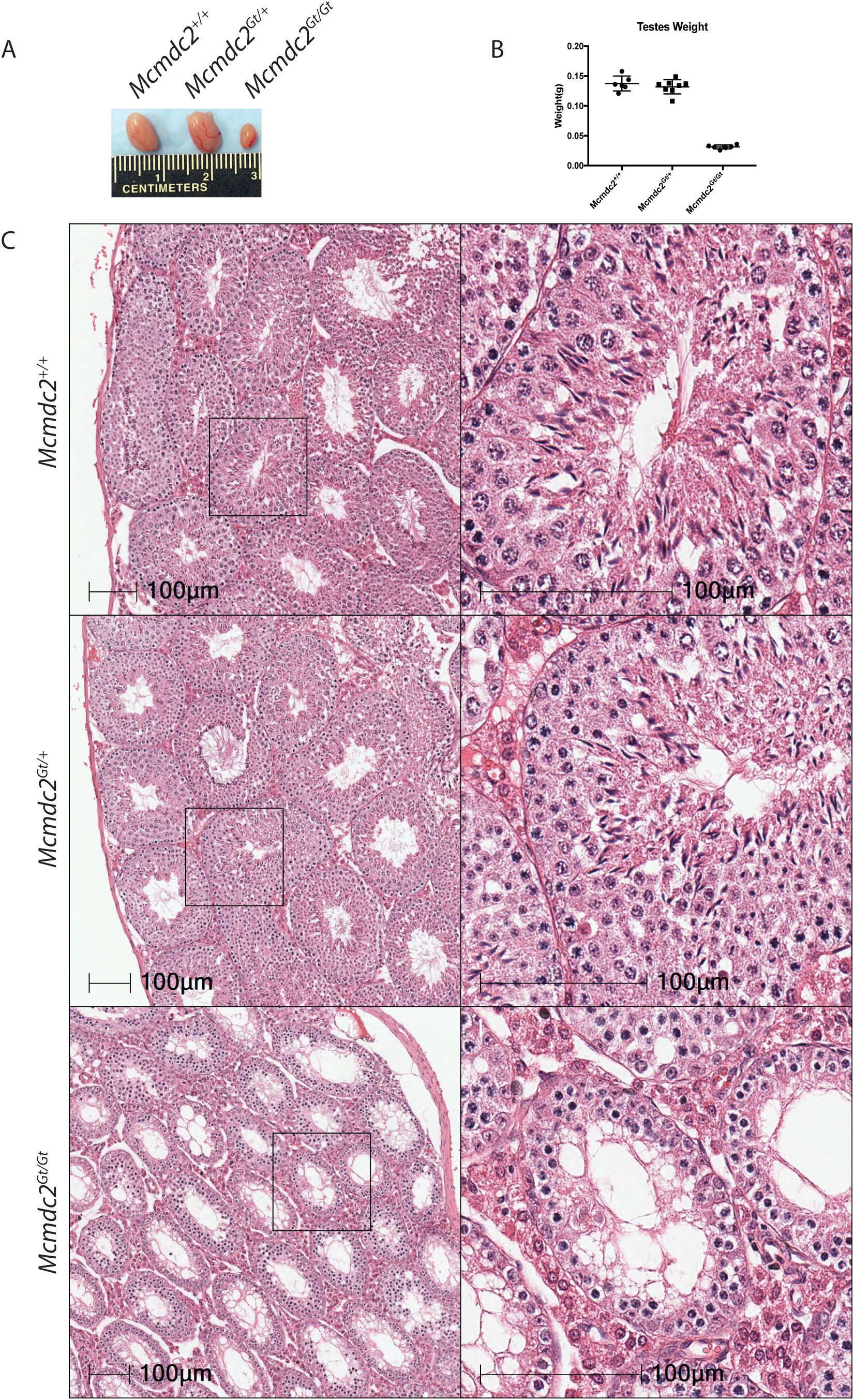
Testes lacking *Mcmdc2* are significantly smaller and lack postmeiotic spermatids. A) Testes from 12-week old littermates of the indicated genotypes. B) Testis weights from *Mcmdc2*^*+/+*^(N=3), *Mcmdc2*^*Gt/+*^(N=4), and *Mcmdc2*^*Gt/Gt*^ (N=4) animals. C) H&E of testes sections from 12-week old animals of the indicated genotypes. Higher magnification of individual tubules are presented in the insets.

**Figure 2.**
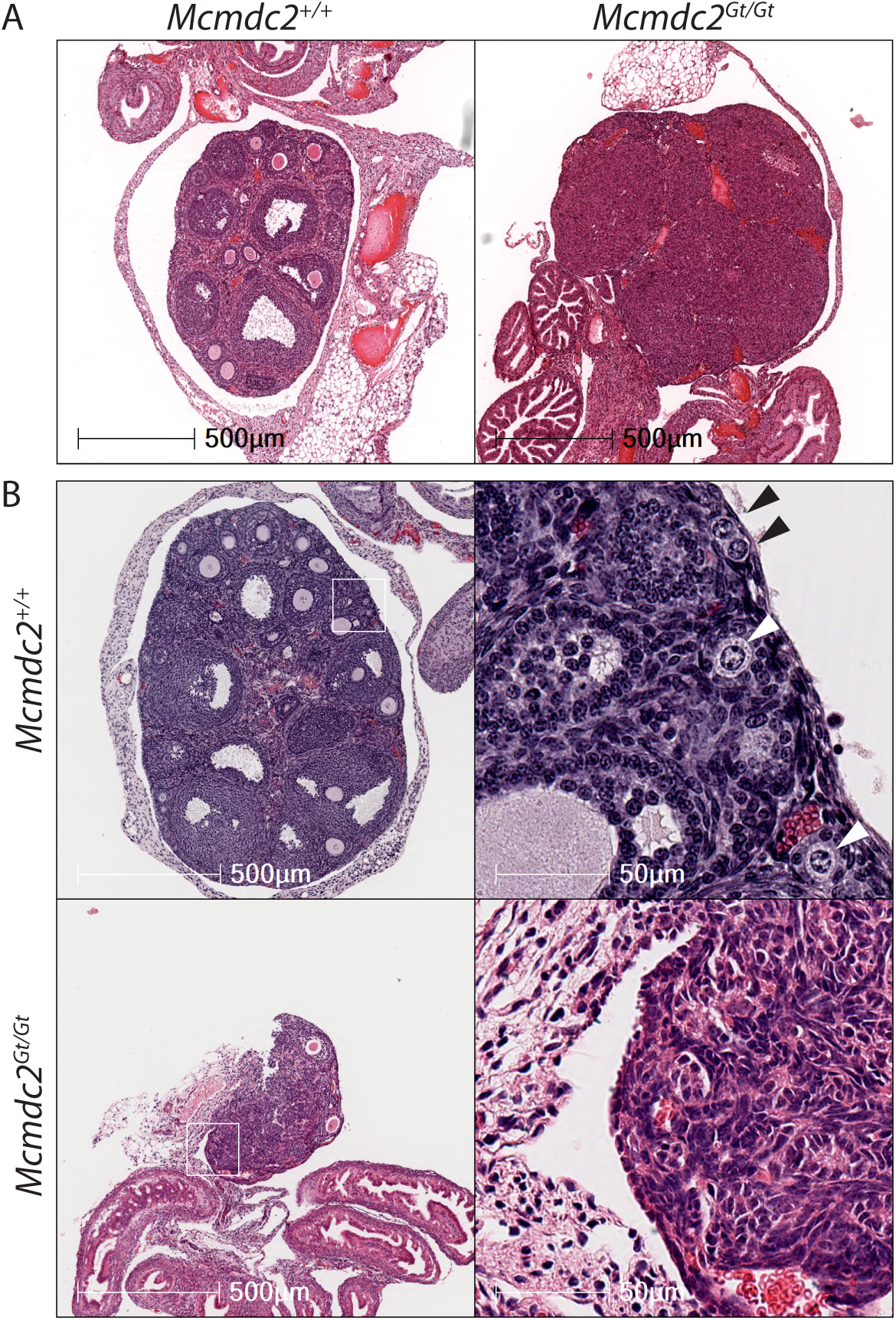
*Mcmdc2^Gt/Gt^* ovarian histology reveal lack of oocytes. A) H&E staining of adult females demonstrates the absence of follicles (and thus, oocytes) in *Mcmdc2*^*Gt/Gt*^ ovaries. B) H&E staining of 24 day old wild-type or *Mcmdc2*^*Gt/Gt*^ females. Wild-type ovaries contain abundant primordial follicles (black arrows) and primary follicles (white arrows), while no primordial follicles are apparent in the *Mcmdc2*^*Gt/Gt*^ ovaries. *Mcmdc2*^*Gt/Gt*^ ovaries also appear smaller than wild-type.

Given the apparent arrest of spermatogenesis in meiotic prophase I, we next evaluated meiotic progression by immunocytological analysis of meiotic chromosomes, using markers of developmental progression, homolog synapsis, and recombination. In WT 19 day postpartum (dpp) mice, spermatocytes produced in the first wave of spermatogenesis would normally have progressed through pachynema, the stage at which homologous chromosomes are fully synapsed. WT pachytene spermatocytes display diffuse nuclear staining of H1t (a marker of mid-late pachynema) (Meistrich *et al.* 1985), and fully synapsed autosomes as indicated by SYCP1 (a synaptonemal complex (SC) central element protein). *Mcmdc2*^*Gt/Gt*^ mutant spermatocytes lacked H1t staining, indicating a block to developmental progression prior to mid pachynema (Fig. 3A,E), and *Mcmdc2*^*Gt/Gt*^ chromosomes remained in a zygotene-like state devoid of appreciable homolog synapsis (Fig. 3B).

**Figure 3.**
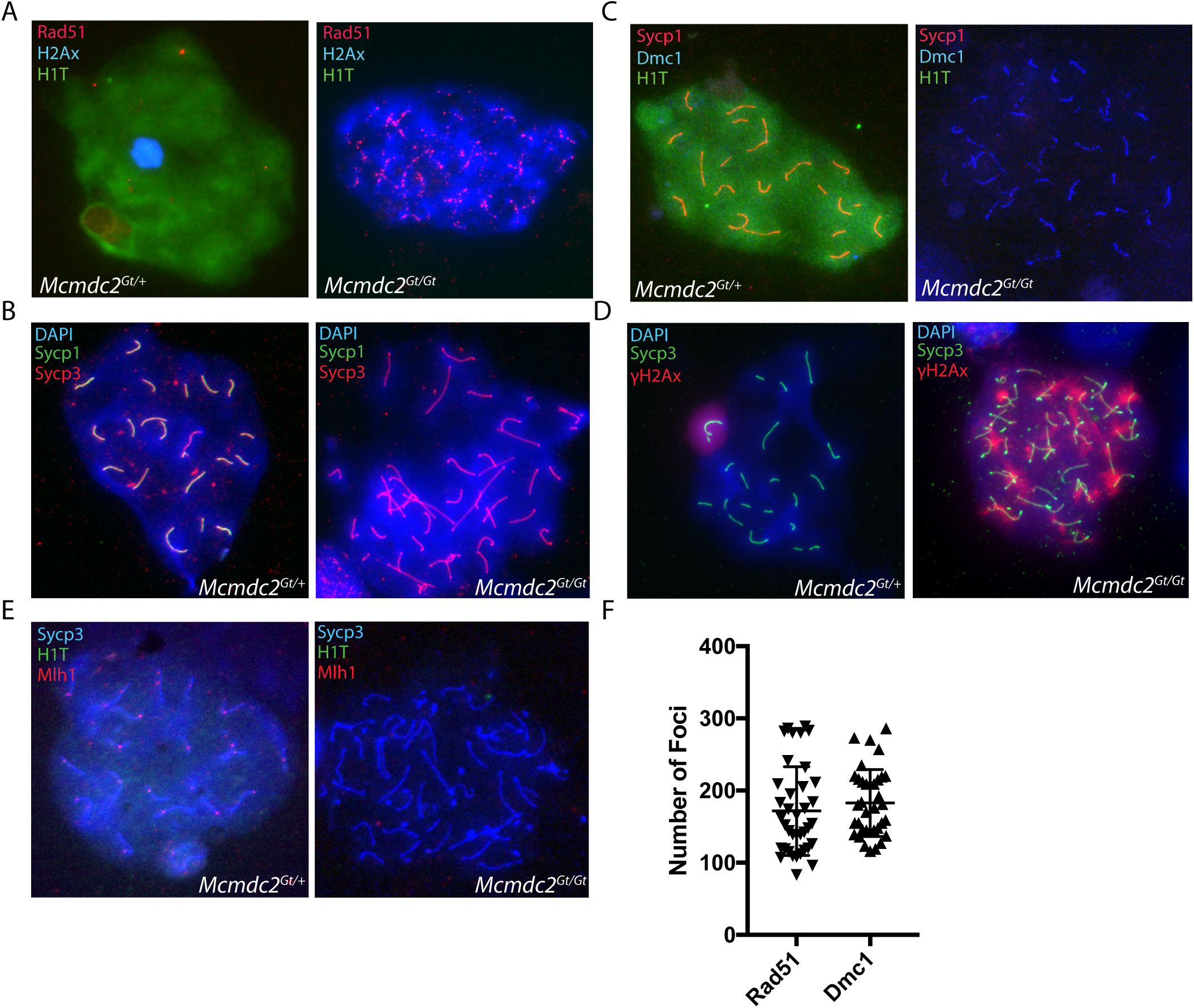
*Mcmdc2*^*Gt/Gt*^ spermatocytes undergo meiotic arrest and are deficient for double strand break repair. Immunofluorescent images of male meiotic spreads from 19 day old *Mcmdc2^Gt/+^* and *Mcmdc2*^*Gt/Gt*^ animals were stained with the indicated antibodies. A) *Mcmdc2*^*Gt/Gt*^ mutants exhibit abundant γH2AX staining, failure to progress to pachynema (absence of H1t), and persistent RAD51 foci. B) *Mcmdc2*^*Gt/Gt*^ mutant chromosomes fail to synapse and form mature SC. C) *Mcmdc2*^*Gt/Gt*^ mutants exhibit persistent DMC1 foci, lack of progression to pachynema, and asynapsis. D) *Mcmdc2*^*Gt/Gt*^ mutants have abundant γH2AX, which marks both asynapsis and unrepaired DSBs. E) *Mcmdc2*^*Gt/Gt*^ spermatocyets lack of MLH1 foci, which mark COs in normal mice. F) Quantification of RAD51 and DMC1 foci in late zygonema-like *Mcmdc2*^*Gt/Gt*^ spread preparations. Each data point represents a single meiotic spread (DMC1, N=36; RAD51, N=38).

A common cause of failed homolog synapsis is defective recombination. Recombination is initiated in leptonema by the formation of several hundred DSBs by the SPO11 protein. As part of a DNA damage response, these breaks trigger ATR-mediated phosphorylation of histone H2AX (gH2AX) on chromatin (Turner *et al.* 2004). The recA homologs RAD51 and DMC1 load onto resected ends of DSBs, forming cytologically visible foci that can be visualized by immunolabeling. These nucleoprotein filaments drive interchromosomal recombination, which is necessary for homolog pairing and synapsis in mice. As Prophase I proceeds, these foci are localized to SC cores, and gradually disappear until DSB repair is complete in pachynema. Whereas these processes appeared normal in heterozygous spermatocytes, there were gross defects in the mutants. First, there were persistent RAD51 and DMC1 along unsynapsed chromosomes axes (Fig. 3C,D). *Mcmdc2*^*Gt/Gt*^ chromosome spreads contained abnormally high levels of persistent RAD51 (171 ± 9.9 foci per spread on average) and DMC1 (182 ± 7.7 foci per spread) indicating a lack of normal DSB repair (Fig. 3F).

In mouse spermatocytes, meiotic prophase I arrest is linked to failure of meiotic sex chromosome inactivation (MSCI). Normally, the X and Y chromosomes are transcriptionally silenced and sequestered in the XY body, a structure that is marked by concentrated γH2Ax and DNA damage response proteins. Disruption of the XY body, which can occur by redirection of silencing machinery to unsynapsed autosomes in a process called MSUC (Meiotic silencing of unsynapsed chromatin), allows expression of Y-linked genes that arrest spermatocyte development (Royo *et al.* 2010). Consistent with the observed lack of DSB repair and asynapsis, *Mcmdc2*^*Gt/Gt*^ spermatocytes had abnormal XY bodies, and exhibited abundant γH2Ax staining over unsynapsed autosomes, which is an indicative of not only persistent DNA-damage, but also MSUC (Fig. 3A,D). These results, along with the histological analysis, indicate MCMDC2 acts early in meiosis to facilitate DNA repair by homologous recombination, and its absence results in meiotic prophase I arrest.

In normal meiocytes, a subset of the meiotic DSBs (~10%) are repaired and resolved as crossovers. In pachynema, CO sites are marked by MLH1 and normally there is at least 1 focus present per chromosome pair. In diplonema, the chromosomes undergo desynapsis and remain attached at sites of COs which are visible as chiasmata. As expected from the gross defect in DSB repair and global synapsis failure that prevents progress through pachynema, no MLH1 foci were detectible in mutant spermatocytes as are present on H1t-positive heterozygous pachytene chromosomes (Fig. 3E).

Our observations show that mouse MCMDC2 is required for both NCO and CO homologous recombination repair of SPO11-induced meiotic DSBs, and in the absence of recombination, synapsis fails to occur. Because DMC1 and RAD51 load onto meiotic chromosomes, we conclude that MCMDC2 functions after DSBs are resected, but before the DMC1/RAD51 filaments invade homologous chromosomes to form stable recombination intermediates in a manner that would drive pairing. It isn’t known if MCMDC2 functions in conjunction with MCM8 (*Drosophila* rec ortholog), but interestingly, the meiotic phenotypes of *Mcm8^-/-^* mutant spermatocytes are quite similar to those of *Mcmdc2*^*Gt/Gt*^ (Lutzmann *et al.* 2012). However, MCM8 has additional functions in S-phase and other tissues; in mitotically growing cells, MCM8 and MCM9 are needed to recruit RAD51 to DSBs and facilitate homologous recombination (Lutzmann *et al.* 2012; Nishimura *et al.* 2012; Park *et al.* 2013). Since MCMDC2 expression is almost exclusive to meiotic cells, any possible functional relationship with MCM8 would be unique to meiosis, and is unlikely related to recruiting RAD51 to SPO11-induced DSBs.

In addition to the similarity with *Mcm8* mutants, *Mcmdc2*^*Gt/Gt*^ phenocopies *Msh4* and *Msh5* knockout mice in that there is defective DSB repair and asynapsis despite loading of RAD51/DMC1 onto DSBs (Kneitz *et al.* 2000; Her *et al.* 2001; Reynolds *et al.* 2013). It is possible that MCMDC2 participates in promoting MSH4/5 function or other members of the ZMM complex that stabilize D-loops formed by invasion of single stranded ends into homologous sequences (Manhart and Alani 2016). Such a scenario would link to the hypothesis that the mei-MCM complex in *Drosophila* adapted to serve the function(s) of MSH4/5 (Kohl *et al.* 2012). The obvious phenotypic distinction between the mouse *Mcm8, Msh4, Msh5* and *Mcmdc2* mutants compared to mei-MCM mutants in *Drosophila* is that in the latter, oocytes survive and can complete meiosis. This is because meiotic DSBs are repaired in the fly mutants (mainly by NCO recombination), and thus activation of the DNA damage checkpoint, which occurs in mutants deficient for recombinational such as the RAD51 ortholog *spn-A*, does not occur (Staeva-Vieira *et al.* 2003) (Ghabrial and Schupbach 1999). The failure of mouse meiocytes to repair DSBs in the *Mcm8, Msh4, Msh5* and *Mcmdc2* mutants may be attributable to an inability to sufficiently stabilize recombination intermediates (following strand invasion) to a degree that would drive pairing and synapsis nucleation. Because interhomolog synapsis occurs independently of recombination in flies, the SC may stabilize recombination intermediates sufficiently to permit DSB resolution in the absence of mei-MCMs. At the moment, there are no proteins known to interact with MCMDC2, and the lack of a suitable available antibody precludes studies of its localization or the purification of complexes in which it may function. Such information will be crucial for understanding the biochemical function of MCMDC2 in mice, and the similarities and distinctions compared to mei-MCMs in *Drosophila*.

## Materials and Methods

### Generation of *Mcmdc2^Gt^* animals

All animals use was conducted under a protocol to JCS approved by Cornell’s Institutional Animal Use and Care Committee. *Mcmdc2*^*Gt*^ ES cells were obtained from EuMMCR (European mutant mouse cell repository) *Mcmdc2^tm4a(EUCOMM)Hmgu^* (*Mcmdc2*^*Gt*^) Founder animals were genotyped by PCR and crossed for 3-4 generations into B6(Cg)-*Tyrc-^2J/^J* B6/2J. To remove the floxed neomycin, *Mcmdc2^tm4a^* females were crossed to a *Stra8*-Cre male (Sadate-Ngatchou *et al.* 2008) and the offspring genotyped for *Cre* and *Mcmdc2*^*Gt*^. The neomycin deleted animals were then backcrossed 2-3 generation into B6. There were no phenotypic differences between neomycin(+) versus neomycin(-) animals. Genotyping primers used are listed in Table S1.

### RT-qPCR

Total RNA was isolated from tissue using E.Z.N.A kit (Omega Biotek) according to manufacturers’ instructions. cDNA was made from 500ug of total RNA using qScript cDNA mix (Quanta), and qPCR carried out using iTaq (Bio-Rad). All RT-qPCR data was normalized to *Gapdh*. Primer sequences are listed in Table S1.

### Histology and LacZ staining

For H&E staining, testes and ovaries were dissected and fixed in Bouin’s for 6 to 12 hours. Tissues were then washed in 70% ethanol for 2-3 days prior to being paraffin embedded. Sections of 4uM thickness were stained with Harris Hematoxylin and eosin (H&E). Slides were digitized using a Leica Scanscope CS2 and 20X lens. Images were analysed using ImageScope and Halo software. For LacZ, tissues was fixed in 4% paraformaldehyde in PBS overnight at 4°C. Tissues were washed with PBS and placed in 30% sucrose/PBS overnight at 4°C. The fixed tissue was then flash frozen in OCT and 10uM sections cut on a cryostat. LacZ staining was carried out as described (Nagy *et al.* 2007).

### Meiotic chromosome surface spreads and immunofluorescence

Isolated testes with the tunica removed were minced into small (2-3mm) pieces in DMEM. Meiocytes were hypotonically swollen in in a sucrose solution on slides. Cells were lysed with 0.1% triton and 0.02% SDS and 2% paraformaldehyde. Slides were washed and stored at -80°C until stained. For staining, slides were brought to room temperature and washed once with PBS+0.1% Triton X-100. Slides were blocked for 40 minutes at RT with 1X PBS+0.3% Triton X-100 containing 5% normal goat serum. Primary antibodies were diluted into 1xPBS/1%BSA/0.3% Triton X-100 and incubated overnight at 37°C in a humidified chamber. Antibodies and dilutions used include: mouse anti-γH2AX Ser 139 (1:1000 Millipore), rabbit anti-Rad51 (1:250 Millipore), rabbit anti-SYCP1 (1:600 Abcam), rabbit anti-SYCP3 (1:600 Abcam), mouse anti-SYCP3 (1:600 Abcam), rabbit anti-MLH1 (1:50 Millipore), mouse anti-DMC1 (1:100 Abcam), and guinea pig anti-H1T (1:1000 gift from M.A.Handel). Secondary antibodies were used at 1:1000 in antibody dilution buffer and included: goat anti-rabbit Alexa 488, goat anti-rabbit Alexa 594, goat anti-mouse Alexa 488/594, and goat anti-guinea pig Alexa 488/594. Images were taken using an Olympus microscope with 63X lens and CCD camera. Foci were quantified using Fiji (Schindelin *et al.* 2012) and analyzed in Prism 7 (Graphpad).

---

**Table S1.**
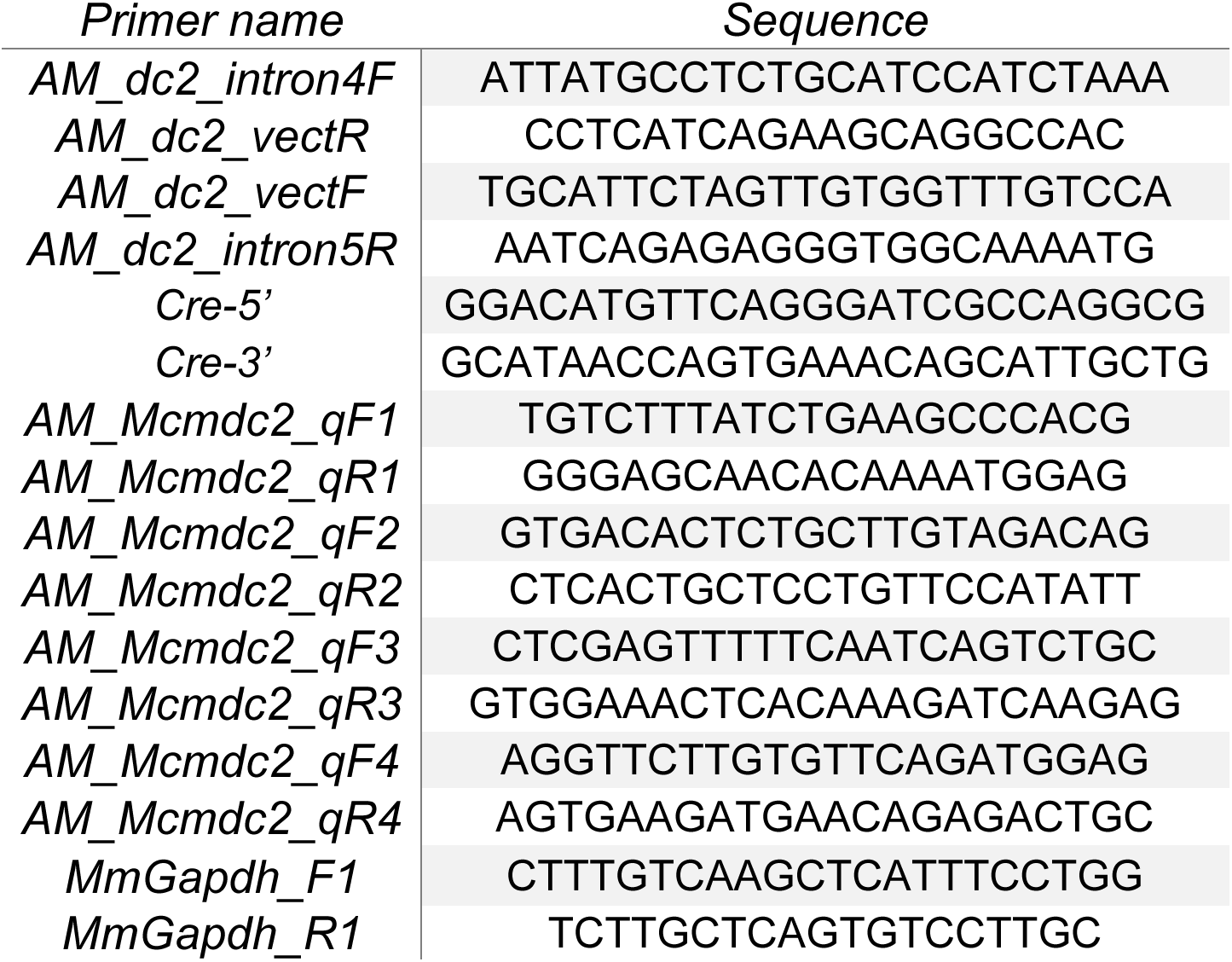
PCR Primers used in study.

**Figure S1.**
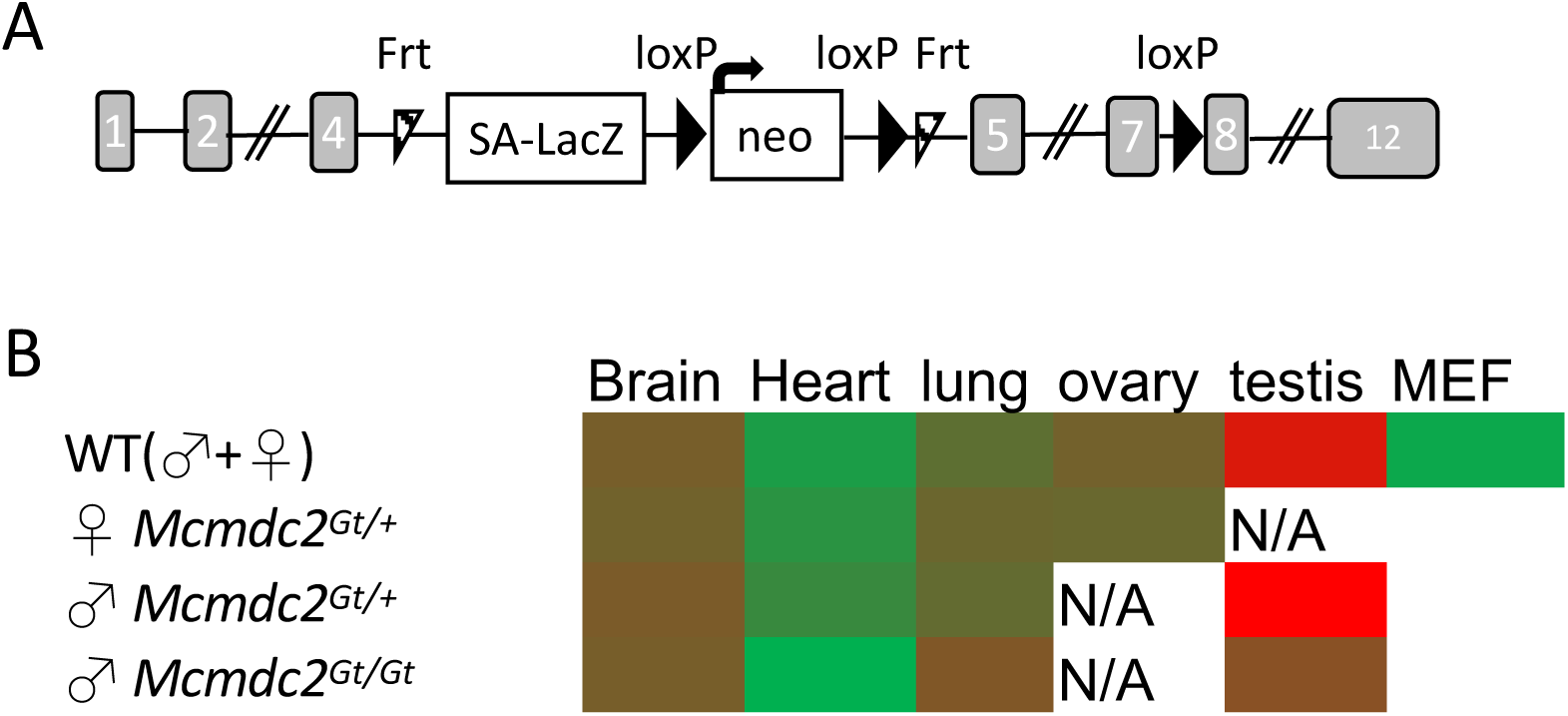
*Mcmdc2* gene trap and expression in mouse. A.)Schematic of ES gene trap vector used in the production of *Mcmdc2^Gt/+^* animals. B.) Heatmap of RT-qPCR results for *Mcmdc2* expression in various tissues and cells. Total RNA was isolated from the indicated tissues and used in qPCR with primers against 3 exons of *Mcmdc2*. Green indicates low or absent expression, red is highest. Ovary expression is likely low as oocytes undergo meiosis embryonically and adult ovaries were examined.

**Figure S2.**
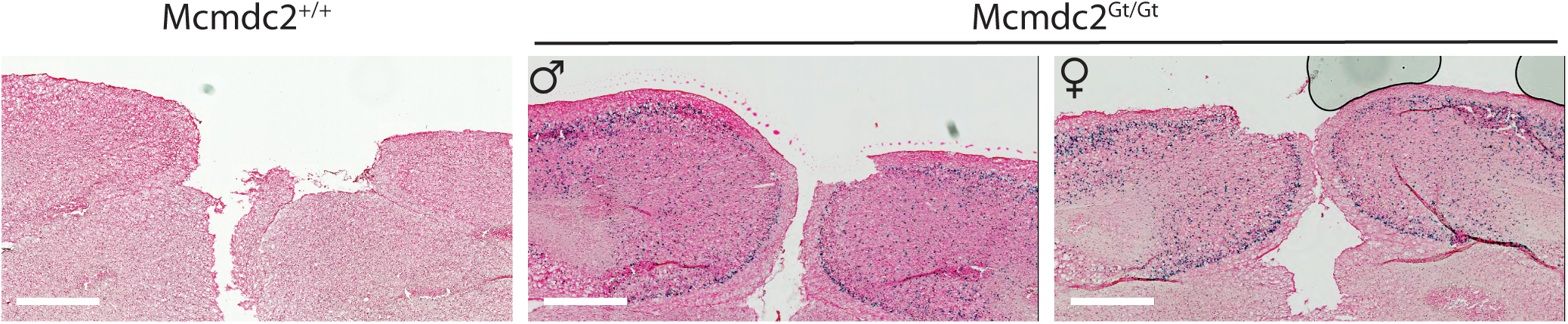
LacZ staining confirms *Mcmdc2*^*Gt*^ expression in brain. To verify expression of the gene-trap in tissues with known Mcmdc2 expression, brains from wild-type and *Mcmdc2*^*Gt/Gt*^ animals were stained for lacZ activity. Both male and female *Mcmdc2*^*Gt/Gt*^ animals exhibited a lacZ staining pattern similar to that seen for *Mcmdc2* mRNA in the Allen Brain Atlas (http://mouse.brain-map.org/experiment/show?id=70227866).

